# SARS-CoV-2 Can Infect Human Embryos

**DOI:** 10.1101/2021.01.21.427501

**Authors:** Mauricio Montano, Andrea R. Victor, Darren K. Griffin, Tommy Duong, Nathalie Bolduc, Andrew Farmer, Vidur Garg, Anna-Katerina Hadjantonakis, Alison Coates, Frank L. Barnes, Christo G. Zouves, Warner C. Greene, Manuel Viotti

## Abstract

The spread of SARS-CoV-2 has led to a devastating pandemic, with infections resulting in a range of symptoms collectively known as COVID-19. The full repertoire of human tissues and organs susceptible to infection is an area of active investigation, and some studies have implicated the reproductive system. The effects of COVID-19 on human reproduction remain poorly understood, and particularly the impact on early embryogenesis and establishment of a pregnancy are not known. In this work, we explore the susceptibility of early human embryos to SARS-CoV-2 infection. We note that ACE2 and TMPRSS2, two canonical cell entry factors for SARS-CoV-2, are co-expressed in cells of the trophectoderm in blastocyst-stage preimplantation embryos. Using fluorescent reporter virions pseudotyped with Spike (S) glycoprotein from SARS-CoV-2, we observe robust infection of trophectoderm cells, and this permissiveness could be attenuated with blocking antibodies targeting S or ACE2. When exposing human blastocysts to the live, fully infectious SARS-CoV-2, we detected cases of infection that compromised embryo health. Therefore, we identify a new human target tissue for SARS-CoV-2 with potential medical implications for reproductive health during the COVID-19 pandemic and its aftermath.

## Introduction

Coronavirus disease 2019 (COVID-19) has emerged as an unexpected and devastating pandemic upending life around the globe [1]. Entry of severe acute respiratory syndrome coronavirus 2 (SARS-CoV-2), the infectious agent responsible for COVID-19 [2], requires interactions of its surface glycoprotein Spike (S) with two entry factors on the target cell including the angiotensin-converting enzyme 2 (ACE2) receptor bound by S and a priming cleavage of S by the serine protease TMPRSS2 [3, 4]. These binding and processing steps promote virion fusion at the plasma membrane [5], or alternatively subsequent endocytosis of virions and fusion within the late endosome [6].

The full repertoire of cell and tissue types that SARS-CoV-2 can infect is now being better defined [1, 7]. Expression of SARS-CoV-2 cell entry factors has been described in a wide assortment of human cells [7], and in COVID-19 patients, SARS-CoV-2 virus has been detected in various organs [8–10]. Infected individuals can exhibit a range of symptoms spanning beyond lung-related problems, to include disease in the intestine, heart, kidney, vasculature, and liver [8,10–14]. Within the female and male reproductive systems, expression of SARS-CoV-2 entry factors is present in cells of the ovaries, uterus, vagina, testis, and prostate [7,15–18]. Some studies showed detectable virus in the semen of infected males [19, 20] and vaginal secretions of infected females [21].

The possibility of vertical transmission of SARS-CoV-2 to embryos during or shortly after fertilization is concerning, but is contingent on cells of the embryo being permissive to the virus. Here, we explore whether the factors required for SARS-CoV-2 entry are present and functional in cells of the preimplantation embryo, and using S pseudotyped reporter viruses and the fully infectious version of the virus, we assess if SARS-CoV-2 can infect human embryos.

## Materials

### Ethical Approval

The study was conducted in adherence with the Declaration of Helsinki. Ethical approval for this project was obtained through the IRB of the Zouves Foundation for Reproductive Medicine (OHRP IRB00011505) and the UCSF Human Gamete, Embryo and Stem Cell Research (GESCR) Committee (Study # 21-34579). Furthermore, the UCSF IRB exempted the study from full review because it does not involve human subjects as defined by federal regulation 45 CFR 46.102(e) of the U.S. Department of Health & Human Services.

### Human Embryos

All embryos used in this study were surplus samples from fertility treatment and *in vitro* fertilization, donated strictly for research by signed informed consent. Embryos were generated using standard industry procedures as previously described [22], vitrified and cryopreserved until thaw and experimental use. For the RNA-seq experiment, embryos were of various ethnic backgrounds and comprised a mix of euploid and aneuploid samples based on evaluation by preimplantation genetic testing for aneuploidy (PGT-A) [23] (Supplementary Table S1). For the pseudotyped virion infection experiments, embryos were either untested or assessed by PGT-A, and included a mix of embryos classified as euploid, mosaic, and aneuploid (samples with aneuploidies in chromosomes X or 21, respectively encoding *ACE2* and *TMPRSS2*, were excluded). For the live virus infection experiments, embryos were used that had not been previously tested by PGT-A. All embryo experiments included a range of initial developmental stages spanning from early to hatched blastocysts.

### RNA-seq and Expression Analysis

Trophectoderm biopsies containing 5-10 cells from blastocyst-stage embryos (n=24) were processed for RNA-seq using a commercial kit (Takara Bio, USA, SMART-Seq v4 Ultra Low Input RNA Kit for Sequencing) following the user manual. The resulting cDNAs were converted to libraries using a Nextera XT kit (Illumina, USA), with a modified protocol according to SMART-Seq v4 user manual. These libraries were pooled and sequenced using a NextSeq 550 instrument (Illumina, USA) with a MidOutput cartridge at 2x75 cycles. The sequencing reads in Fastq files were down-sampled to 6M total reads, aligned to the human genome assembly (hg38), and the number of transcripts per million (TPM) was determined using the CLC Genomics Workbench 12 (Qiagen, USA). Results were mined for expression of factors implicated in SARS-CoV-2 infection. Violin plots were prepared with PlotsOfData [24].

### Immunostaining

Blastocysts were immersed in fixation buffer containing 4% paraformaldehyde (EMS, USA, no. 15710) and 10% fetal bovine serum (FBS; Seradigm, USA, 1500-050) in phosphate-buffered saline (PBS; Corning, USA, MT21040CM) for 10 minutes (min) at room temperature (rt), followed by three 1-min washes at rt in PBS with 10% FBS. To inhibit non-specific protein interactions, embryos were cultured in 2% horse serum (Sigma, USA, H0146) diluted in PBS (blocking solution) followed by addition of 0.1% saponin (Sigma, USA, S7900) for gentle permeabilization. After 1 h incubation at rt, primary antibodies diluted in blocking solution were added and incubated overnight at 4 °C. Embryos were then washed three times for 5 min each in PBS at rt prior to incubation with secondary antibodies. Secondary antibodies diluted in blocking solution were applied for 1 h at 4 °C. Embryos were then washed twice for 5 min each in PBS and subsequently incubated with 5 µg/ml Hoechst 33342 (Invitrogen, USA) in PBS for 5 min to stain the nuclei. Finally, embryos were washed twice for 5 min each in PBS prior to mounting for imaging. The following primary antibodies were used: goat anti-ACE2 (R&D Systems, USA, AF933, 1:100), mouse anti-TMPRSS2 (Developmental Hybridoma Bank, USA, P5H9-A3, 3.2 µg/ml). The following secondary Alexa Fluor-conjugated antibodies (Invitrogen, USA) were used at a dilution of 1:500: donkey anti-goat Alexa Fluor 568 (A10042), donkey anti-mouse Alexa Fluor 488 (A21202). DNA was visualized using Hoechst 33342. For all immunofluorescence experiments, five independent experiments were performed and analyzed.

### Pseudotyped Virion Preparation

For production of HIV-1 NL-43ΔEnv-eGFP SARS CoV-2 S pseudotyped virus particles, 293T cells were plated at 3.75 x10^6^ cells in a T175 flask. 24 h post plating the cells were transfected using a PEI transfection reagent (Sigma, USA). 90 µg of PEI, 30 µg of HIV-1 NL-4ΔEnv-eGFP expression vector DNA (NIH AIDS Reagent Program, USA) and 3.5 µg of pCAGGS SARS CoV-2 S Glycoprotein expression vector DNA (NR52310, BEI, USA) in a total of 10 ml of Opti-MEM media (Invitrogen). The day following transfection the media was changed to DMEM10 complete media and samples were placed at 37 °C and 5% CO_2_ for 48 h. At 48 h, the supernatants were harvested, filtered through 0.22 µm Steriflip filters (EMD, Millipore, USA) and concentrated by ultracentrifugation for 1.5 h at 4 °C at 25K rpm. After concentration, the supernatant was removed and pellets containing virus particle pellets were resuspended in cold 1xPBS containing 1% FBS, and aliquots stored at -80 °C. For production of control virus particles not expressing the SARS CoV-2 S glycoprotein (Bald), the same procedure was followed except the pCAGGS SARS CoV-2 S vector DNA was omitted from the transfection. SARS and MERS pseudotyped virus particles were produced using the same procedure, substituting the SARS CoV-2 S expression vector with either pcDNA3.1(+) SARS S or pcDNA3.1(+) MERS S.

For production of VSVΔG SARS CoV-2 S pseudotyped virus particles, 293T cells were plated at 1.8 x10^6^ cells in a T175 flask. 24 h post plating, the cells were transfected by PEI transfection reagent (Sigma, USA) with 90 µg of PEI, 30 µg of pCAGGS SARS CoV-2 S Glycoprotein expression vector DNA (NR52310, BEI, USA) in a total of 10 mL of Opti-MEM media (Invitrogen, USA). One day after transfection the media was removed, the cells were washed with 1xPBS and DMEM10 complete media was added. Once the media was changed the cells were infected with VSVΔG VSVg virus (Sandia, USA) at an MOI of 1 or higher. The infection media was changed after 4 h, the cells were washed with 1xPBS and DMEM10 supplemented with 20% anti-VSVg hybridoma supernatant (ATCC, USA, CRL-2700). At 24 h the supernatant was harvested, filtered by 0.22 µm Steriflip filter (EMD, Millipore, USA) and then concentrated by ultracentrifugation for 1.5 h at 4 °C and 25K rpm. Supernatant was removed and virus particle pellets were resuspended in cold 1xPBS containing 1% FBS, aliquots were stored at -80 °C. For production of control virus particles not expressing the SARS CoV-2 S glycoprotein (Bald), the same procedure was used but with the omission of the pCAGGS SARS CoV-2 S vector transfection on day 2.

SARS and MERS S pseudotyped virus particles were produced using the same procedures, substituting the SARS CoV-2 S expression vector with either pcDNA3.1(+) SARS S or pcDNA3.1(+) MERS S vectors respectively.

### Pseudotyped Virion Infection Assay

Blastocyst-stage embryos were hatched from zona pellucidas mechanically (except for samples that were already fully hatched), and transferred to flat bottom 96 well plates in 100 µl embryo culture media. Either HIV-1 NL-43ΔEnv-eGFP SARS CoV-2 S pseudotyped virions (100ng/p24), or VSVΔG SARS CoV-2 S pseudotyped virions (MOI=0.1), were added to the embryos. Bald (not expressing S glycoprotein) virions and mock infection conditions were included in each pseudotyping experiment. After the addition of the virions, the embryos were spinoculated with virus by centrifugation at 200 g for 2 h at rt. Upon completion of the spinoculation, an additional 100 µl of embryo culture media was added to each well and the cultures were placed at 37 °C and 5% CO_2_. For the HIV-1 NL-43ΔEnv-eGFP based infections, embryos were monitored for fluorescence at 24-48 h post-spinoculation. For the VSVΔG based infections embryos were monitored for fluorescence at 12-24 h post-spinoculation. Additional controls included the addition of 10 µg of anti-ACE2 antibody (AF933, R&D Systems, USA) that blocks SARS-CoV-2 S binding to ACE2, anti-SARS CoV-2 S Neutralizing antibody (SAD-S35, ACRO, USA) that interacts with the receptor binding domain on Spike that blocks Spike engagement of the ACE2 receptor, or anti-Human IgG Kappa (STAR 127, Bio-Rad, USA) control antibody

### Live SARS-CoV-2 Viral Infection Assay

Viral stocks were prepared using Vero E6 cells using an infectious molecular clone of SARS-CoV-2 expressing an mNeonGreen reporter (icSARS-CoV-2-mNeonGreen) [25]. For the infection assay, blastocyst-stage embryos were placed in 96 well flat bottom plates in 100 µl of embryo culture media. A viral preparation at a concentration of 100 TCID_50_ in culture media was added to the embryos. Control embryos were additionally incubated with either 10ug of anti-ACE2 antibody (AF933, R&D Systems, USA), anti-SARS CoV-2 Spike Neutralizing antibody (SAD-S35, ACRO, USA) or anti-Human IgG Kappa (STAR 127, Bio-Rad, USA) antibody. The plates were spinoculated at 200g for 1 hour at room temperature. Upon completion of the spinoculation an additional 100uLs of embryo culture media was added to each well and the cultures were placed at 37°c and 5% CO_2_. At 12 to 16 hours post-spinoculation the embryos were assessed for infection. All experiments were performed in the Gladstone Institutes ABSL3 facility, adhering to BSL3 protocols.

### Microscopy

Embryos were placed into 35-mm glass-bottom dishes (MatTek, USA). For epifluorescence microscopy, embryos were imaged with an Echo Revolve fluorescent microscope (Echo, USA) or an EVOS M5000 Imaging System (ThermoFisher, USA) employing a LPanFL PH2 20X/0.40 lens, and fluorescence light cube for GFP (470/525 nm) and transmitted light. For confocal microscopy of immunostained embryos, samples were suspended in small quantities of a 4 mg/ml solution of BSA (Sigma, USA) in PBS. Images were acquired using a LSM880 (Zeiss, Germany) laser-scanning confocal microscope, equipped with an oil-immersion Zeiss EC Plan-Neofluar 40x/NA1.3/WD0.17mm. Z-stacks were acquired through whole embryos with an optical section thickness of 1 µm. Fluorescence was excited with a 405-nm laser diode (Hoechst), a 488-nm Argon laser (Alexa Fluor 488), and a 561-nm DPSS laser (Alexa Fluor 568). For confocal microscopy of infected embryos, samples were stained with Hoechst 33342, and images were acquired using an Olympus FV3000RS laser-scanning microscope using a 40X UPLXAPO (NA=0.95). Embryos were simultaneously scanned for Hoechst and GFP using the 405-nm and 488-nm lasers. Rapid Z series were taken through the entire volume of the imaged embryos in 3 µm steps.

## Results

### Embryo Cells Express Genes Required for SARS-CoV-2 Infection

We reasoned that within the preimplantation period of human development, embryos in the blastocyst stage may become vulnerable to SARS-CoV-2 as they lose their protective zona pellucida. This structure counters the threat of many foreign agents [26]. The trophectoderm, which is the precursor of the placenta [27], is located at the surface of the blastocyst and may be an early target for infecting viruses. Hence, we focused our attention on the trophectoderm and evaluated its permissiveness to SARS-CoV-2 infection.

Two prior publications have described *ACE2* and *TMPRSS2* expression in blastocyst-stage preimplantation embryos [7, 28], albeit based on analysis of publicly available RNA-seq datasets from a single ethnic group (East Asian/Chinese) [29–31]. To determine whether this pattern of gene expression is observed in more diverse populations, we performed RNA-seq on a group of 24 human blastocysts from multiple ethnic backgrounds (Supplementary Table S1). *ACE2* transcripts were detected in trophectoderm biopsies comprising 5-10 cells in 23 of 24 embryos (95.8%), and *TMPRSS2* transcripts in biopsies from all 24 embryos (Fig. 1A). The group of tested embryos ranged in developmental stage from early blastocyst to hatched blastocyst, meaning that *ACE2* and *TMPRSS2* transcripts were present throughout this developmental window. We evaluated the expression of 22 additional human genes proposed to be involved in the SARS-CoV-2 life cycle [7] (Fig. 1B). Positive expression was confirmed for genes encoding some putative alternate receptors (*BSG/CD147*, *ANPEP*) but not for others (*CD209*, *CLEC4G*, *CLEC4M*). There was also detectable expression of genes encoding some alternative proteases (*CTSB*, *CTSL*, *TMPRSS4*), but not others (*TMPRSS11A* and *TMPRSS11B*). Transcripts for *DPP4*, encoding the receptor used by MERS-CoV S for entry, were either absent or expressed at very low levels in our samples.

**Figure 1.**
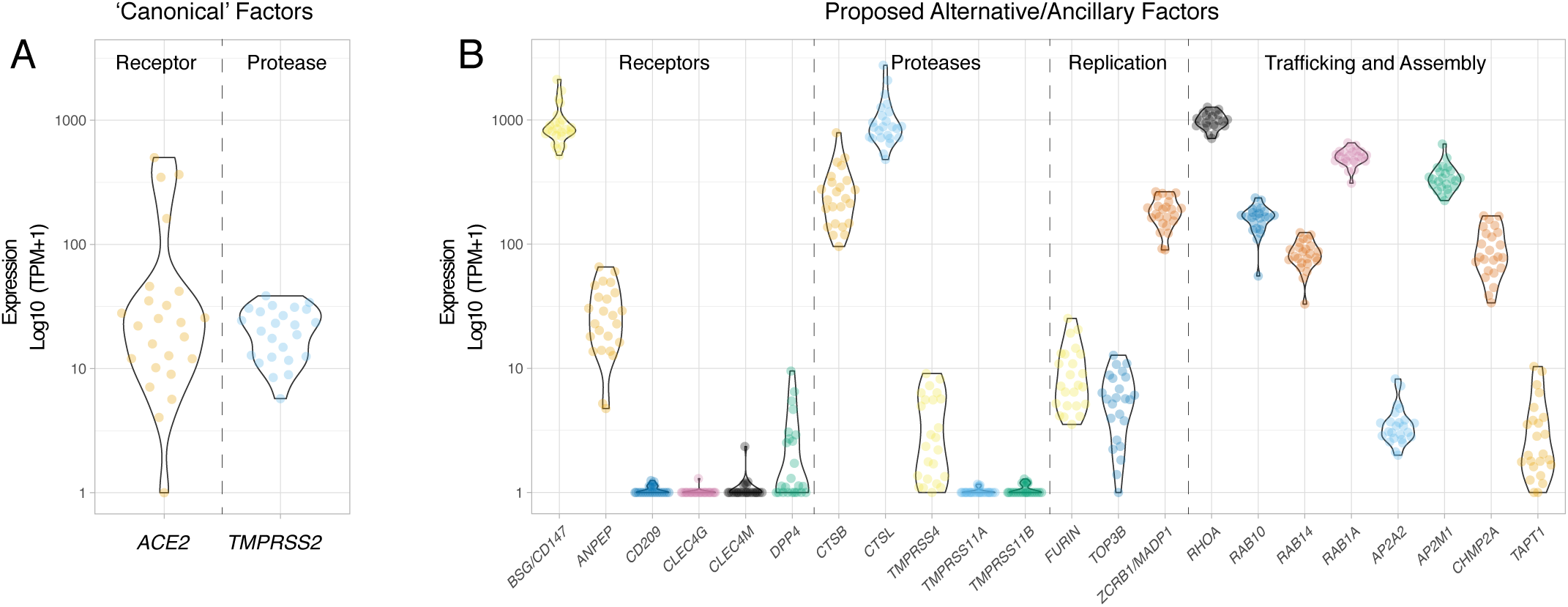
Expression of Genes Involved in SARS-CoV-2 Infection in Embryo Cells. Violin plots showing log10-normalized expression profiles obtained by RNA-seq performed on trophectoderm biopsies of blastocysts. Each data point represents one embryo (n=24). Each trophectoderm biopsy consisted of 5-10 cells. (A) Canonical SARS-CoV-2 entry factors *ACE2* and *TMPRSS2* (B) Proposed alternative/ancillary mediators of SARS-CoV-2 entry, replication, traffic, and assembly.

We noted expression of genes for three factors apparently required for SARS-CoV-2 genome replication (*FURIN*, *TOP3B* and *ZCRB1/MADP1*). Among the genes encoding factors proposed to control trafficking and/or assembly of viral components and which are known to interact with SARS-CoV-2 proteins, *RHOA*, *RAB10*, *RAB14*, *RAB1A*, *AP2M1*, and *CHMP2A* exhibited high levels of expression while *AP2A2* and *TAPT1* were expressed at lower levels. Together, these transcriptomic profiling data indicate that trophectoderm cells express many key factors required for SARS-CoV-2 entry and subsequent replication.

### SARS-CoV-2 Entry Factors Localize to the Membrane of Trophectoderm Cells

To evaluate the presence and localization of entry factors in trophectoderm cells, we performed immunofluorescence and confocal imaging for ACE2 and TMPRSS2 in blastocysts. Both factors were readily detectable in cells of the trophectoderm; ACE2 was enriched on cellular membranes, as evidenced by strong signal at cell-cell junctions (Fig. 2A), while the TMPRSS2 presence was more diffuse, localizing to both cell membranes and within the cytoplasm but not nucleus (Fig. 2B). The inner cell mass (ICM) was not evaluated, since the immunofluorescence protocol was not optimized for penetration of antibodies into deeper cell layers. Based on the localization of ACE2 and TMPRSS2 proteins, at least trophectoderm cells of preimplantation embryos contain the requisite entry factors needed for SARS-CoV-2 infection.

**Figure 2.**
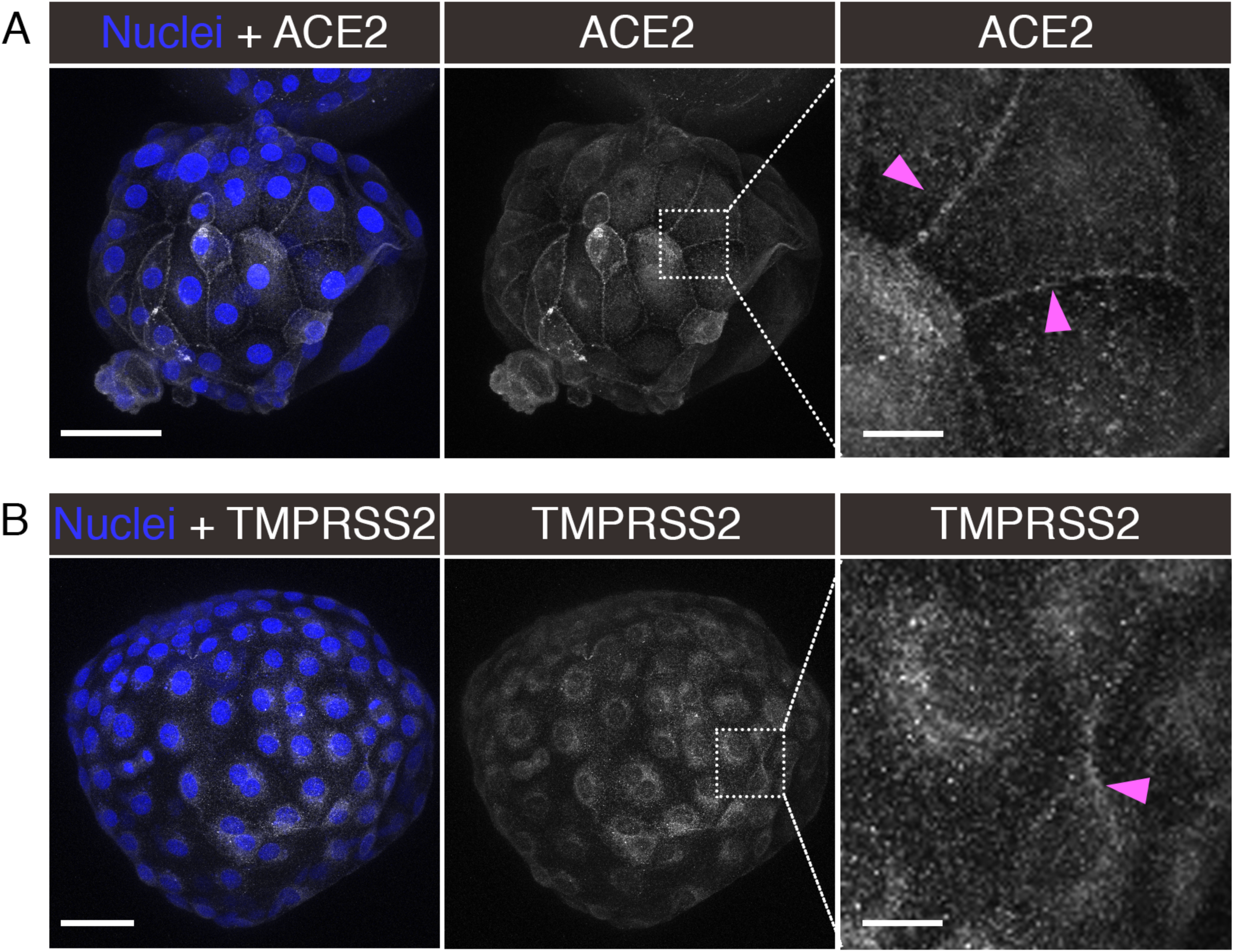
Localization of ACE2 and TMPRSS2 in Embryos. Maximum intensity projections (MIPs) of confocal z-stacks of blastocysts, showing nuclei (blue) and ACE2 or TMPRSS2 (white). Pink arrowheads point to cell membranes. Scale bars represent 50 µm in low magnification panels, and 10 µm in high magnification panels. Representative images from independent biological replicates (n=5 embryos for each factor).

### S-Pseudotyped Virions Utilize ACE2 for Entry Into Trophectoderm Cells

To test whether the standard S protein-mediated SARS-CoV-2 cell-entry process was functional in embryo cells, we evaluated the entry of pseudotyped reporter virions expressing S in zona pellucida-free blastocysts. In the first series of experiments (see summary on Table 1), we used an HIV-ΔEnv virus encoding a green fluorescence protein (GFP) reporter. Embryos exposed to control media or media containing the original non-pseudotyped ‘bald’ reporter virus displayed no fluorescence and appeared healthy 24-48 hours after mock spinoculation (inoculation by centrifugation) (Supplementary Fig. S1). However, when embryos were exposed to the reporter virus pseudotyped with the S protein from SARS-CoV-2, several showed robust GFP signal in numerous trophectoderm cells (Fig. 3A and Supplementary Fig. S1). Treatment of the blastocysts with neutralizing anti-S antibodies markedly decreased GFP fluorescence to a limited number of puncta (Supplementary Fig. S1).

**Figure 3.**
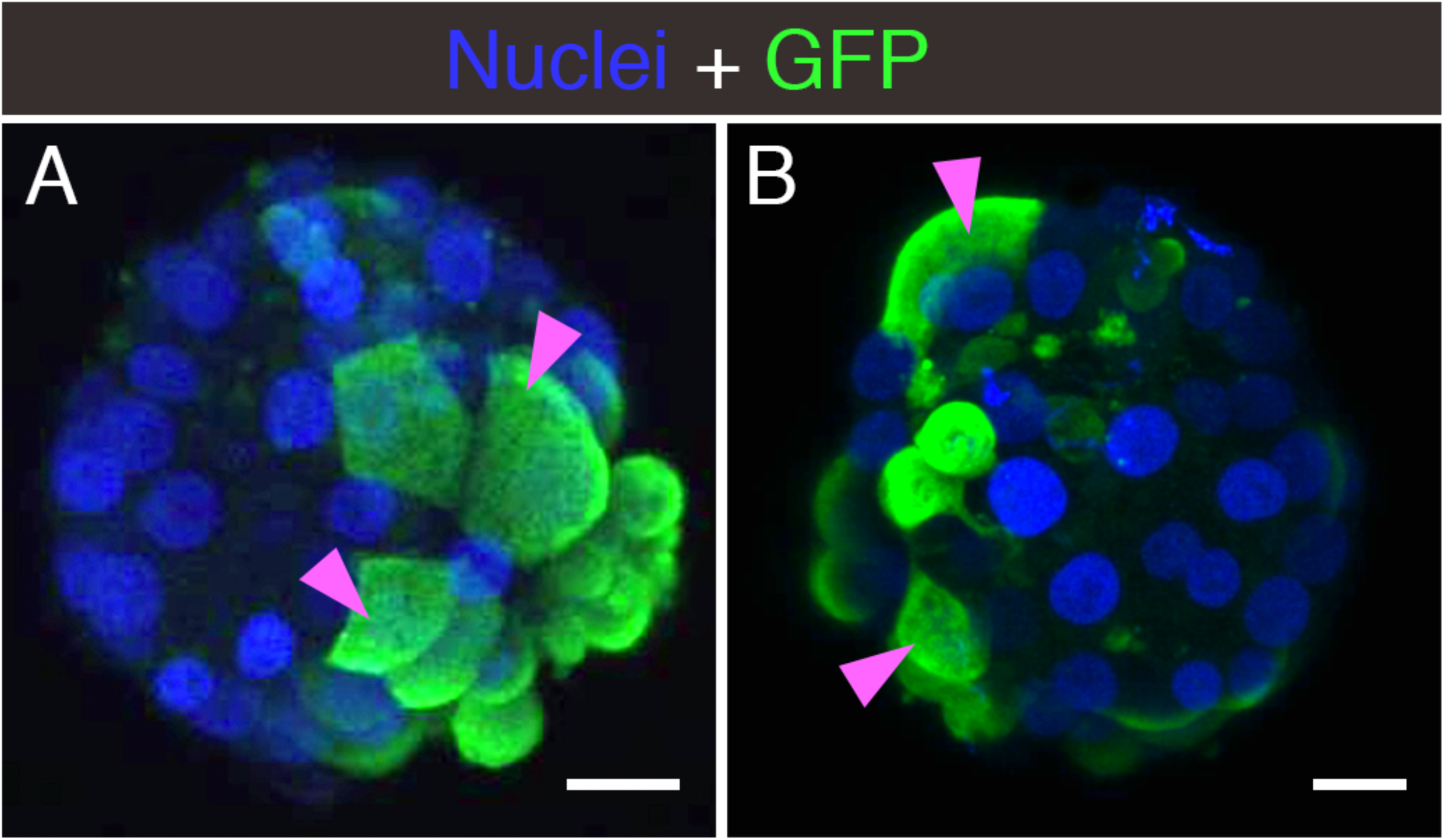
Embryo Infection by Reporter Virions Pseudotyped with the S protein of SARS-CoV-2. Sample confocal MIP images of embryos infected with (A) HIV-based or (B) VSVΔG - based reporter virions pseudotyped with the S protein from SARS-CoV-2. Pink arrowheads point to cells displaying robust signal. Scale bars represent 20 µm. Representative images from independent biological replicates (n=15 embryos with HIV-based virion, n=7 embryos with VSVΔG -based virion).

**Table 1.**
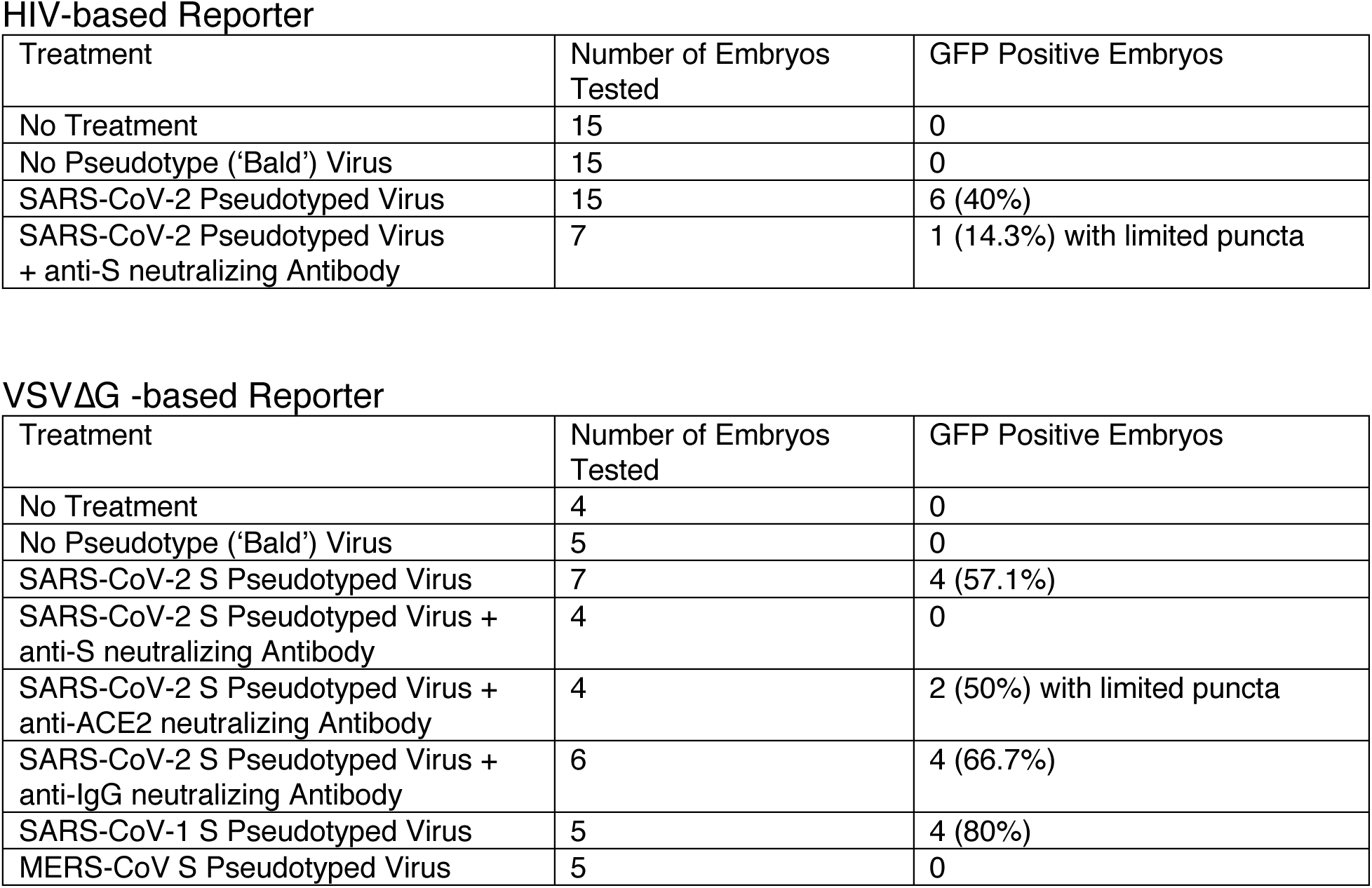
Summary of Experiments Using Pseudotyped Reporter Virions. For each reporter virion (HIV- or VSVΔG -based), the table indicates the experimental condition, the number of embryos used, and the number/percent of infected embryos as evidenced by GFP signal.

HIV-based Reporter
| Treatment | Number of Embryos Tested | GFP Positive Embryos |
| --- | --- | --- |
| No Treatment | 15 | 0 |
| No Pseudotype ('Bald') Virus | 15 | 0 |
| SARS-CoV-2 Pseudotyped Virus | 15 | 6 (40%) |
| SARS-CoV-2 Pseudotyped Virus + anti-S neutralizing Antibody | 7 | 1 (14.3%) with limited puncta |

**Table 1. Summary of Experiments Using Pseudotyped Reporter Virions.**
| Treatment | Number of Embryos Tested | GFP Positive Embryos |
| --- | --- | --- |
| No Treatment | 4 | 0 |
| No Pseudotype ('Bald') Virus | 5 | 0 |
| SARS-CoV-2 S Pseudotyped Virus | 7 | 4 (57.1%) |
| SARS-CoV-2 S Pseudotyped Virus + anti-S neutralizing Antibody | 4 | 0 |
| SARS-CoV-2 S Pseudotyped Virus + anti-ACE2 neutralizing Antibody | 4 | 2 (50%) with limited puncta |
| SARS-CoV-2 S Pseudotyped Virus + anti-IgG neutralizing Antibody | 6 | 4 (66.7%) |
| SARS-CoV-1 S Pseudotyped Virus | 5 | 4 (80%) |
| MERS-CoV S Pseudotyped Virus | 5 | 0 |

In the second experimental series (see summary on Table 1), we used a vesicular stomatitis virus lacking the cell entry factor G glycoprotein (VSVΔG), and encoding GFP as a reporter. No fluorescence was detected when embryos were exposed to control media or media containing the non-pseudotyped ‘bald’ virions . Conversely, several samples exhibited GFP when exposed to the reporter virus pseudotyped with the SARS-CoV-2 S protein (Fig. 3B and Supplementary Fig. S1). Addition of a neutralizing antibody targeting either S or ACE2 strongly reduced GFP expression, while addition of a control non-specific anti-IgG antibody resulted in undiminished GFP expression in several embryos (Supplementary Fig. S1). When the VSVΔG-based reporter virus was pseudotyped with the S protein from SARS-CoV-1, which also utilizes the ACE2 receptor for cell entry, embryos again displayed GFP signal. Conversely, reporter virus pseudotyped with the S protein from MERS-CoV, which depends on the dipeptidyl peptidase 4 (DPP4) receptor for cell entry, produced no GFP signal (Supplementary Fig. S1).

In these experiments using pseudotyped reporter virions, embryos with evidence of infection displayed occasional cell degradation, likely due to expression of native genes in the HIV- and VSV-based reporter virions (Supplementary Fig. S1).

Together, these pseudotyped-virion experiments indicate that trophectoderm cells present in preimplantation embryos are permissive to SARS-CoV-2 entry involving interactions between S and the ACE2 receptor.

### Live SARS-CoV-2 Infects Human Embryos

We next tested the susceptibility of embryos to live SARS-CoV-2 infection. Embryos were exposed to a version of SARS-CoV-2 that, in addition to possessing its full native infectivity, also expresses a fluorescent reporter protein facilitating visualization of infected cells [25]. For this experiment, we did not remove the zona pellucida from blastocysts growing in vitro. As a result, the samples contained a mix of embryos at different stages of blastocyst development, ranging from non-hatched (fully encapsulated by the zona pellucida), to hatching (some cells herniating out of an opening in zona pellucida), to fully hatched (all embryo cells have fully emerged from the zona pellucida).

Some embryos readily displayed evidence of infection when exposed to the virus (Fig. 4A and Table 2), which could be prevented with neutralizing antibodies targeting S or ACE2, but not with a non-specific anti-IgG antibody (Supplementary Fig. S2 and Table 2). Infected cells were predominantly in the trophectoderm, but in some instances, there was also evidence of infection in the ICM (Fig. 4B). Infection was uncommon in non-hatching blastocysts, common in hatching blastocysts, and frequent in fully hatched blastocysts (Fig. 5A). Only one non-hatching blastocyst displayed evidence of infection (Fig. 5B). Some hatching blastocysts only contained infected cells in the herniating (zona pellucida-free) compartment (Fig. 5C). Others had infected cells in both herniating and non-herniating compartments, however cells proximal to the zona opening were more readily infected than cells distal to the zona opening (Fig. 5D-E). We confirmed that experimental groups showing no infection (no virus, anti-S, and anti-ACE2) did not contain a overrepresentation of non-hatching embryos (Supplementary Table S2).

**Figure 4.**
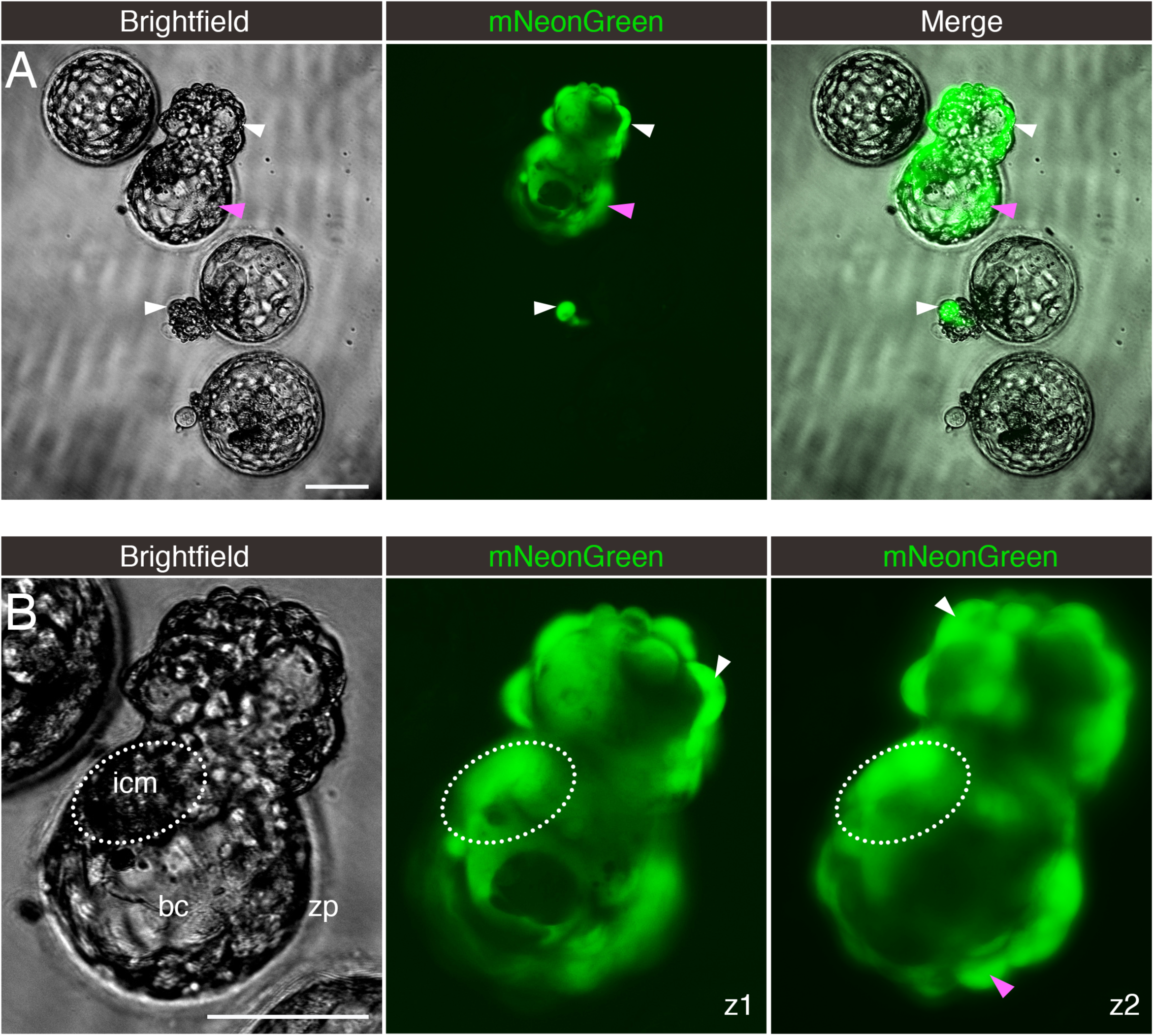
Embryos Exposed to Live SARS-CoV-2 are Susceptible to Infection. Fluorescent signal indicates cells infected with icSARS-CoV-2-mNeonGreen. (A) Four sample blastocysts exposed to the virus. One blastocyst has not initiated hatching and shows no evidence of infection, two blastocysts are at early stages of hatching (one shows evidence of infection in the herniating cells), and one blastocyst is in advanced hatching phase showing high incidence of infected cells. (B) High magnification of blastocyst with high incidence of positive cells, with two panels showing the mNeonGreen channel at different focus depths (z-axis). White arrowheads point to positive herniating cells in hatching blastocysts, pink arrowhead points to positive cells in the zona pellucida compartment of a hatching blastocyst. Representative images from independent biological replicates (n=19 embryos). icm = inner cell mass. bc = blastocoel cavity. zp = zona pellucida. Scale bar represents 100 µm.

**Figure 5.**
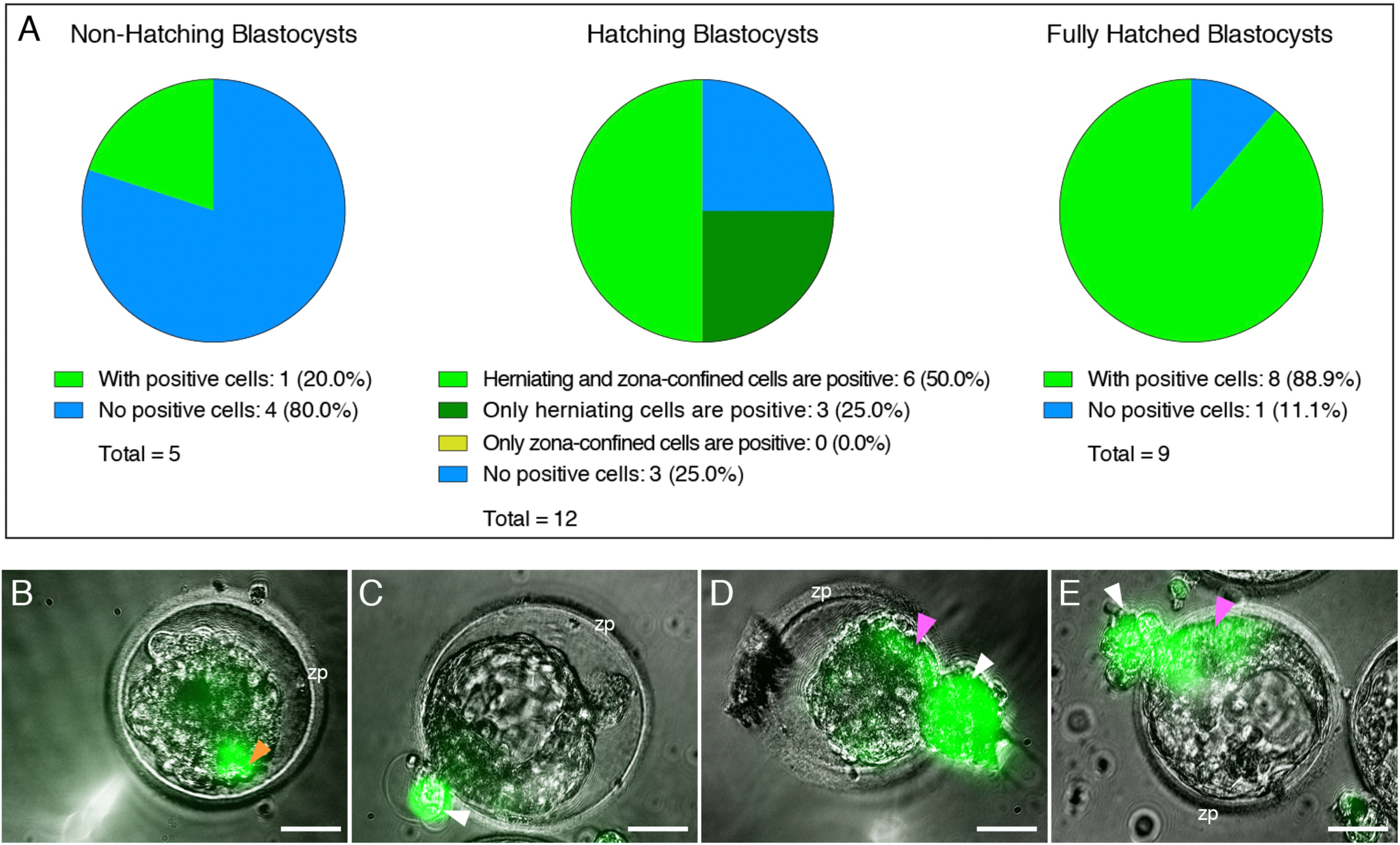
The Zona Pellucida Might Confer Protection Against SARS-CoV-2 Infection. (A) Pie charts indicating the proportion of blastocysts displaying evidence of infection at three separate stages of development (before, during, and after hatching), when exposed to live SARS-CoV-2 expressing a fluorescent reporter (in the presence or absence of control IgG blocking antibody). Non-hatching blastocysts have an intact zona pellucida, hatching blastocysts have some cells herniating out of the zona pellucida opening as well as some cells confined in the zona pellucida compartment, and fully hatched blastocysts have emerged completely out of the zona pellucida. (B) Only example of a non-hatching blastocyst with zona pellucida-encapsulated positive cells (orange arrowhead). (C) Example of a hatching blastocyst with positive cells exclusively in the herniating compartment. (D, E) Examples of hatching blastocysts, with positive cells in the herniating compartment (white arrowheads) as well as in cells proximal to the zona pellucida opening (pink arrowheads). zp = zona pellucida. Scale bars represent 50 µm.

**Table 2.** Summary of Experiments Using Live, Fully-Infectious SARS-CoV-2. The table indicates the experimental condition, the number of embryos used, and the number/percent of infected embryos as evidenced by fluorescent signal.

| Treatment | Number of Embryos Tested | Embryos w/ Fluorescent Cells |
| --- | --- | --- |
| No Treatment | 9 | 0 |
| Virus | 19 | 14 (73.7%) |
| Virus + anti-S neutralizing Antibody | 4 | 0 |
| Virus + anti-ACE2 neutralizing Antibody | 4 | 0 |
| Virus + anti-IgG neutralizing Antibody | 7 | 4 (57.1%) |

These live SARS-CoV-2 virus experiments show that embryos are susceptible to infection, and the zona pellucida might confer protection.

### Infection by SARS-CoV-2 Induces Cytopathicity and Affects Embryo Health

In the course of the SARS-CoV-2 live virus experiments we noted an impact on embryo health. Embryos with evidence of infection (with visible reporter signal) often displayed signs of cell degeneration. This ranged from focal cell degradation, particularly in infected (fluorescent) areas of the embryo, to severely affected embryo health, and at times to embryo collapse and global cell death (Fig. 6A-F). On average, infected embryos exhibited morphologic signs of considerably poorer health and morphology at 12-16 hours post-infection compared to embryos that had gone through the same spinoculation protocol but had not been infected. Out of 18 embryos with evidence of infection, only one appeared in good health, while four displayed focal damage, five exhibited severe cellular degeneration (but were still alive), and eight embryos had completely collapsed and were dead (Fig. 6G). Out of the 25 embryos with no evidence of infection, 16 appeared in good health, while eight showed some cell damage, and one had severe cytopathic changes (likely due to the stress of spinoculation and subsequent culturing) (Fig. 6G).

**Figure 6.**
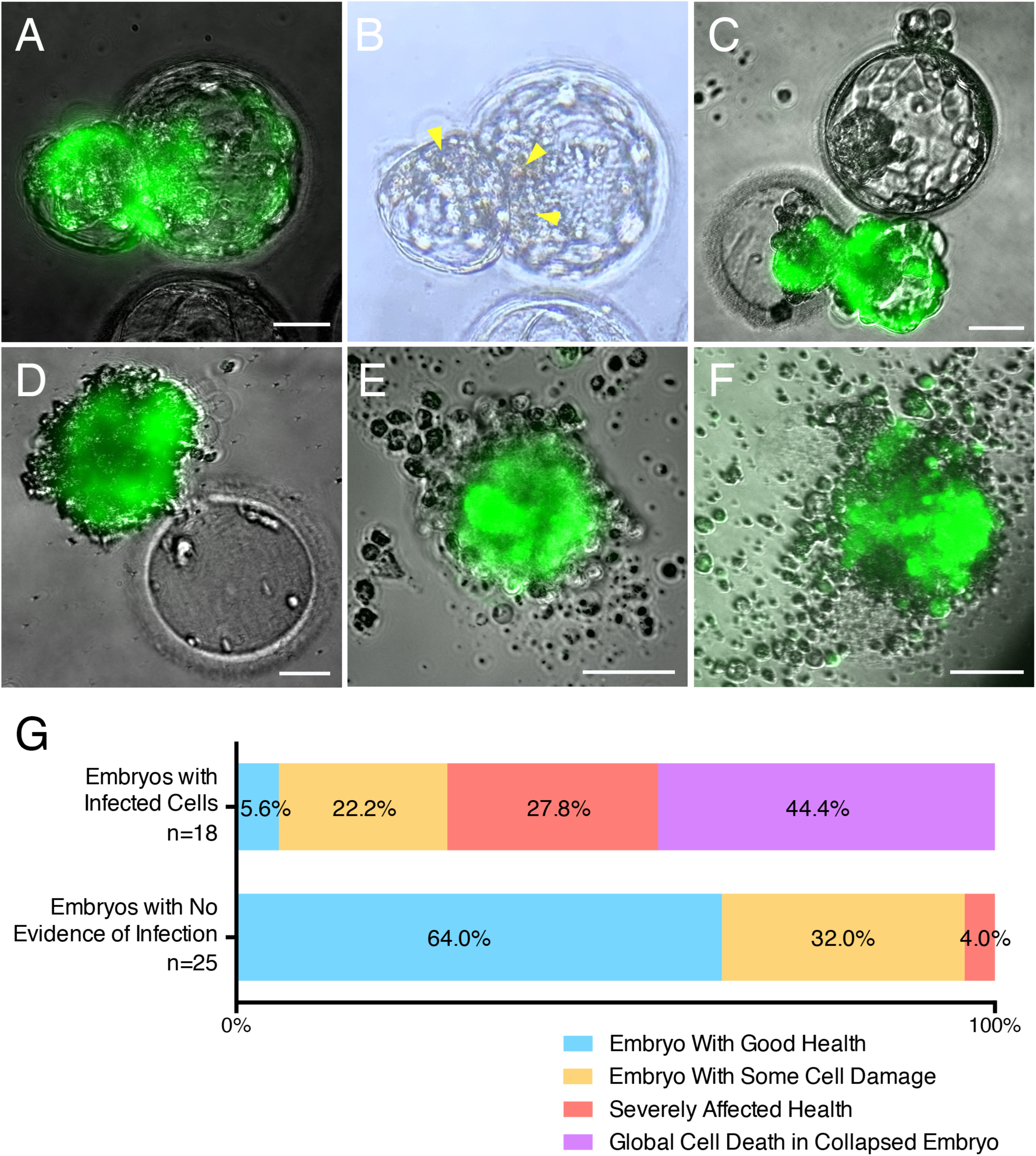
Embryo Health is Negatively Affected by SARS-CoV-2 Infection. Sample images of infected embryos with various degrees of cytopathic effects. (A-B) Embryo with cell damage (yellow arrowheads) in fluorescent (infected) cells in the herniating and zona pellucida compartments. (C) Infected herniating embryo (bottom) with severely affected health, and non-infected embryo (top) with good health. (D) Hatched infected embryo next to empty zona pellucida, displaying global cell death. (E-F) Examples of infected embryos displaying total collapse and death. (G) Summary of embryo health 12-16 hours after exposure to SARS-CoV-2. Top bar includes all embryos with evidence of infection (fluorescent cells). Bottom bar includes all embryos with no evidence of infection (no fluorescent cells). Scale bars represent 50 µm.

These findings suggest a cytopathic effect of infection, which can at times lead to embryo demise.

## Discussion

This study shows that some cells within the human embryo 1) express and correctly localize factors required for SARS-CoV-2 entry, 2) are permissive to viral entry via the S-ACE2 ligand-receptor system, 3) are susceptible to live virus infection, which can produce cytopathic effects, and 5) may be protected against SARS-CoV-2 infection by the zona pellucida. Together, these findings confirm that human embryos at the blastocyst stage prior to implantation are susceptible to SARS-CoV-2 infection, which can impact their viability.

The RNA-seq experiments show that embryos from multiple ethnic backgrounds express the canonical entry factor genes *ACE2* and *TMPRSS2* in trophectoderm cells (conceptually the most vulnerable cell lineage to viral infection as these cells form the outer cell layer of the embryo and ultimately give rise to the placenta). These findings confirm and extend prior reports of *ACE2* and *TRMPRSS2* embryonic expression in a single ethnic group (East Asian/China) [29–31]. Membrane localization of the SARS-CoV-2 receptor ACE2 observed on trophectoderm cells confirms and extends a previous immunofluorescence microscopy study employing a different commercially-available anti-ACE2 antibody [32]. In addition, we further detect TMPRSS2 protein localization in these same cells. TMPRSS2 cleavage of S primes the S2 component for effective virion fusion [3, 4].

The experiments using reporter virions corroborate involvement of S and ACE2. Only virions pseudotyped with the SARS-COV-2 S protein could infect cells of the embryo. Addition of blocking antibodies reactive with either S or ACE2 reduces infection by the pseudotyped virions, implicating a functional interplay between S and ACE2 for virion entry. Embryos also displayed evidence of infection by reporter virions pseudotyped with the S protein from SARS-CoV-1 (which similarly uses ACE2 for entry) but not with the S protein from MERS-CoV (which uses DPP4 for entry). These findings further support the conclusion that ACE2 functions as an effective SARS-CoV-2 receptor in trophectoderm cells.

Finally, using live SARS-CoV-2, we confirm that trophectoderm cells of the embryos are susceptible to viral infection. Infection may also extend to the ICM (the precursor of the fetus). Of note, not all embryos exposed to SARS-CoV-2 showed evidence of infection and when infection occurred, not all of the cells uniformly expressed the viral reporter. Infection is likely dependent on numerous factors, including: grade of an embryo at the time of exposure, timing of infection, and effective titers of the virus (viral copy numbers to which cells are exposed). Our goal was not to achieve maximum infection efficiency, but rather to determine whether cells of the embryo are at all permissive to infection. The observation that the zona pellucida confers protection from infection was reinforced in studies showing preferential infection of cells herniating from the zona during hatching. Embryos infected with live virus displayed evidence of viral cytopathic effects ranging from focal changes to death of the entire embryo likely determined by the overall levels of infection achieved in each embryo.

The finding that preimplantation embryos are susceptible to SARS-CoV-2 infection raises the possibility of viral transmission from either the mother or father to the developing embryo. Vertical transmission of SARS-CoV-2 between pregnant mothers and fetuses has been reported [33–38], albeit considerably later in pregnancy. Of note, these studies implicate the placenta, which develops from the trophectoderm, as the principal site of viral infection and subsequent transmission [35,38–40]. It is estimated that placental infection occurs in 21% of COVID-19 pregnancies, while 2% of neonates show SARS-CoV-2 positivity [41]. A placenta infected by SARS-CoV-2 often shows signs of injury characterized by trophoblast necrosis [42, 43]. Studies analyzing the delivered placentas of SARS-CoV-2 positive pregnant women revealed widespread and diffuse trophoblastic damage sufficient to undermine fetal health [38, 44].

Could vertical transmission of SARS-CoV-2 occur whereby preimplantation embryos are infected? Numerous studies have failed to identify virus in male and female reproductive organs of infected patients [45–47]. However, a more recent study detected the virus in vaginal swabs of two out of 35 women with COVID-19 [21], and two groups have reported virus in semen of infected men [19, 20]. For example, a study by Li and colleagues found that six of 38 male COVID-19 patients had detectable levels of SARS-CoV-2 in their semen [19]. Virus in vaginal fluid or semen could potentially infect a newly conceived embryo prior to implantation, potentially compromising establishment or maintenance of the pregnancy. In view of this, our data highlights the need for expectant mothers or couples considering pregnancy to fully vaccinate to help prevent SARS-CoV-2 infection that could jeopardize their health and their pregnancy [48–50].

In comparison to embryos from natural conceptions, embryos generated by assisted reproductive technologies (ART) for treatment of infertility, such as *in vitro* fertilization (IVF), face additional risks of potential viral exposure. These include exposure to virus shed by asymptomatically infected medical and laboratory personnel during various procedures (handling of gametes, assisted conception, embryo culture, biopsy, vitrification, and thawing), or by infected patients (for example, a male patient contaminating a sample vial during sperm collection, via regular exhalation). The data presented here should inform best practices in the ART clinic during the COVID-19 pandemic, further building on existing recommendations for safety and risk mitigation [51–55].

SARS-CoV-2 infection in pregnant women is associated with increased risk of miscarriage, prematurity, and impaired fetal growth [56]. Such adverse fetal outcomes have mainly been attributed to COVID-19-related complications in pregnant patients [56], but could reflect infection of the placenta or fetus during pregnancy [33-36,40,44]. The present study raises the possibility of vertical transmission during preimplantation stages. Of note, the zona pellucida appears to protect the embryo from infection (likely starting at the zygote stage) but inevitably the blastocyst becomes exposed at hatching, which is a necessary event prior to implantation into the maternal endometrium. Given the trophectoderm’s central role in implantation, compromised health of trophectoderm cells due to SARS-CoV-2 infection could altogether impede establishment of a pregnancy. The same result (no pregnancy) would inevitably occur in the the case of complete embryo collapse, as we observed in a substantial proportion of infected blastocysts. Alternatively, lasting detrimental effects on the trophectoderm-derived placenta could affect the clinical outcome of an established pregnancy by increasing the risk of spontaneous abortion, intrauterine growth restriction (IUGR), or stillbirth, as has been noted in instances of placental infection [38, 44].

Noting that our transcriptomic analysis revealed RNA presence of various factors associated with downstream steps of the SARS-CoV-2 viral life cycle, such as genome replication, trafficking and assembly [7], the possibility of trophectoderm cells infecting surrounding tissues (maternal or fetal) after additional viral shedding cannot be excluded. Our observation that the ICM might itself be vulnerable to infection further highlights this risk. Ultimately, population effects of the COVID-19 pandemic on fertility may become apparent when epidemiological data on pregnancies and birth rates become more readily available.

A weakness of the study is that it primarily focuses on one embryonic tissue, namely the trophectoderm: 1) the RNA-seq experiment specifically probed trophectoderm cells, 2) The immunostaining experiments used a protocol optimized for cell membrane-bound factors on the surface of the embryo (precluding analysis of deeper cell layers), 3) The pseudotyped virion experiments made use of non-replicative agents, meaning that even with entry into the outer cell layer, subsequent infection of deeper layers was not possible. Only the live, fully infectious (and replicative) SARS-CoV-2 experiment was informative regarding potential infection of the ICM. It should be noted however, that by forming the outside layer of the blastocyst, the trophectoderm constitutes the primary site of exposure to viral infection. This means we assessed the likeliest first step of an embryo infection by SARS-CoV-2. A further caveat of the study is that it exclusively used *in vitro*-generated embryos, and the findings might not be entirely applicable to preimplantation embryos derived from natural conceptions.

Future experiments will need to define the presence/absence of virus in the setting of natural conceptions and IVF treatment around embryos, to better understand the risks and take appropriate measures. Also, future work should continue addressing the direct consequences of infection on individual cells, embryo viability, and pregnancy outcomes.

In summary, our finding that preimplantation embryos are susceptible to SARS-CoV-2 infection *in vitro* raises the potential vulnerability of these embryos *in vivo*. Additionally, the data presented here should prompt careful review of procedures surrounding assisted reproduction during the COVID-19 pandemic and its aftermath.

## Author contributions

MV initiated the research question and together with WCG supervised all aspects of the study. The study design was devised by MV and WCG, with input from MM, ARV, FLB, CGZ, DKG, AKH, and AF. Embryology experiments were performed by ARV, virology and imaging experiments were performed by MM, RNA-seq experiments were performed by MV, TD, and NB, immunostaining and fluorescent imaging were performed by MV and VG. MV wrote the first version of the manuscript with input from WCG. All authors contributed to further writing and critical revisions of the manuscript and all authors approved the final version of the manuscript.

## Acknowledgements

We thank Dr. Julie Laliberte, Chelsae Madrid, Dr. Francesca Spinella, and the embryology staff at Zouves Fertility Center for facilitating this study. We also thank Dr. Melanie Ott and members of the Ott lab for their assistance and for facilitating the live virus experiments in the Gladstone BSL-3 facility.

## Data availability

All data generated or analyzed during this study are included in this published article (and its Supplementary Information files).

## Supplementary Tables

**Supplementary Table S1.** Features of Embryos used in RNA-seq Experiment. Chromosomal status (euploid/aneuploid) as determined by PGT-A and background (ethnicity/region) of embryos tested.

| Embryo Number | Ploidy | Affected Chromosomes | Ethnicity/Region |
| --- | --- | --- | --- |
| 1 | Euploid | - | Central American (Mexico) |
| 2 | Euploid | - | South Asian (India) |
| 3 | Euploid | - | African/Mediterranean |
| 4 | Euploid | - | European |
| 5 | Euploid | - | African/Mediterranean |
| 6 | Euploid | - | Middle Eastern (Jewish) |
| 7 | Euploid | - | European |
| 8 | Euploid | - | African/Mediterranean |
| 9 | Aneuploid | +21 | East Asian (China) |
| 10 | Aneuploid | +7,-21 | Central American/European |
| 11 | Aneuploid | -15 | Central American/European |
| 12 | Aneuploid | +5 | European |
| 13 | Aneuploid | -17 | South Asian (India) |
| 14 | Aneuploid | -21,-22 | Central American/European |
| 15 | Aneuploid | +7,+22 | East Asian (China) |
| 16 | Aneuploid | +4 | East Asian (Japan)/European |
| 17 | Aneuploid | +16,+20 | East Asian (China) |
| 18 | Aneuploid | -15,+21 | South-/East Asian (Vietnam/China) |
| 19 | Aneuploid | +6 | South East Asian (Indonesia)/European |
| 20 | Aneuploid | +15,+16 | East Asian (China) |
| 21 | Aneuploid | +11,+12,-22 | East Asian (China) |
| 22 | Aneuploid | -2 | East-/South Asian (China/India) |
| 23 | Aneuploid | +16 | South Asian (India) |
| 24 | Aneuploid | +15 | European |

**Supplementary Table S2.**
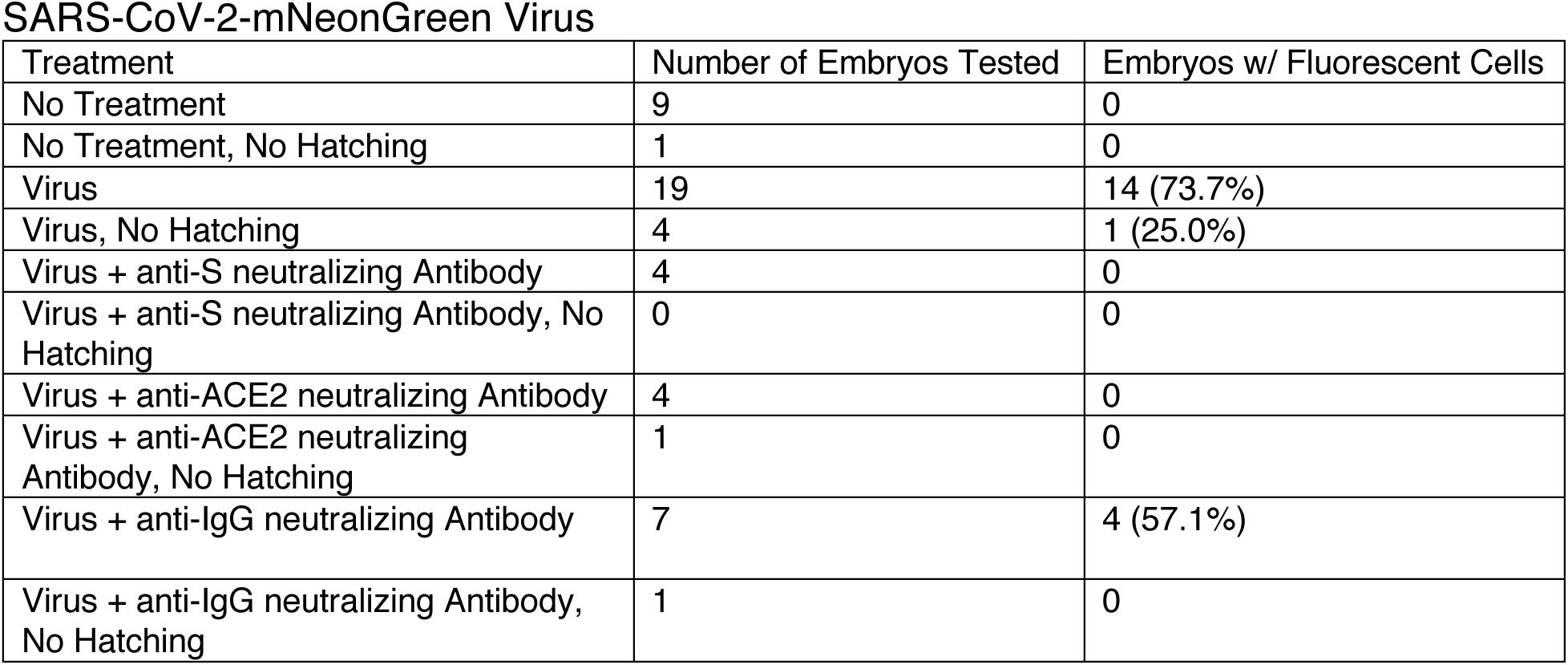
Representation of Non-Hatching Embryos in Experimental Groups of the Live SARS-CoV-2 Experiments. Breakdown of embryos by their (non-)hatching status.

**Supplementary Figure S1.**
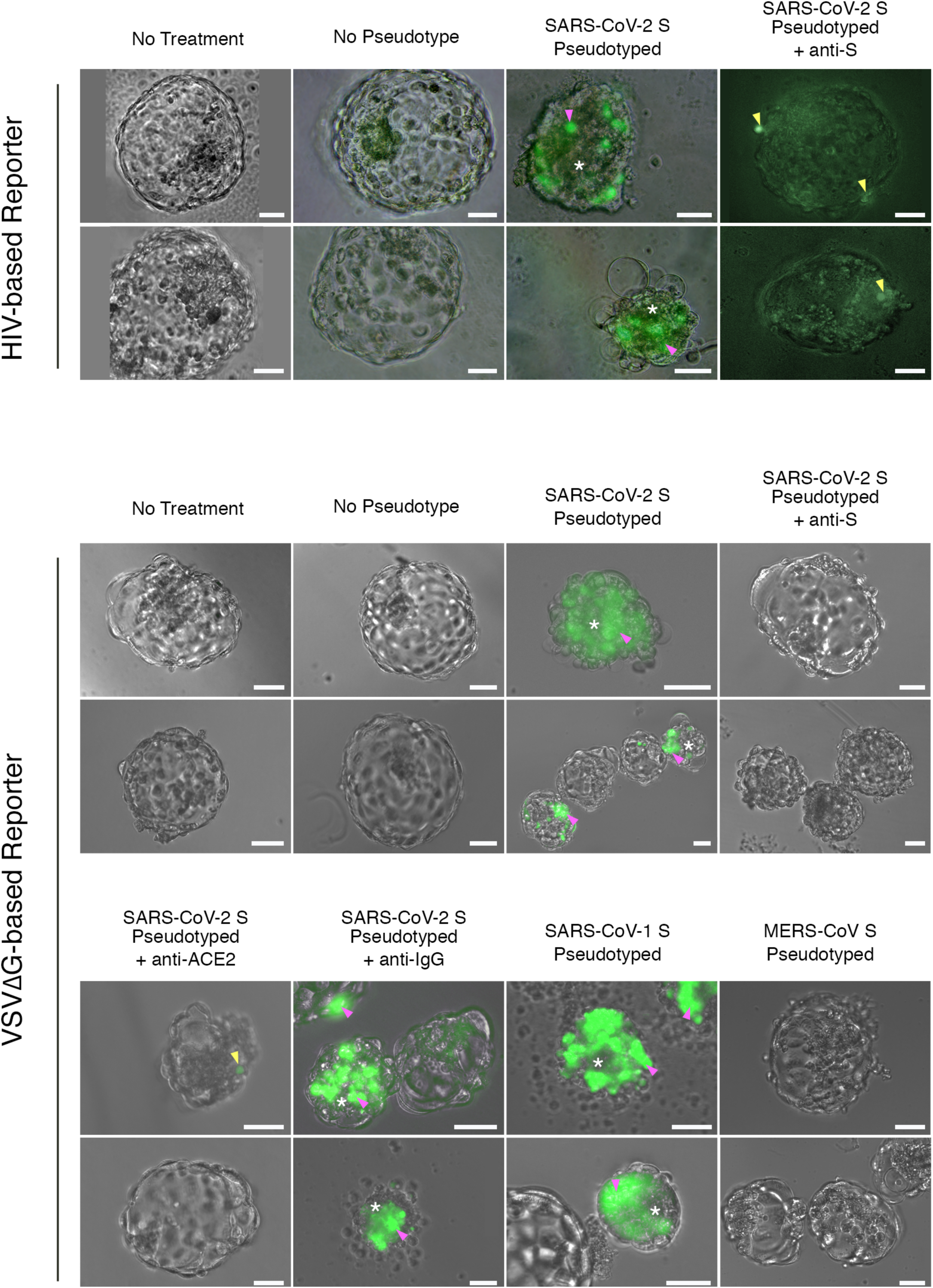
Reporter Virion Experiments Indicate Entry Into Cells of the Embryo Occurs Via S and ACE2. Sample images from GFP reporter virion experiments, displaying merged brightfield with epifluorescence signal. Top set shows results from HIV-based virus, bottom set shows results from the VSVΔG-based virus. Two representative images are shown per condition. The sample size of each condition is indicated in Table 1. Pink arrowheads point to cells displaying robust GFP signal, yellow arrowheads point to punctate GFP signal, and white asterisks indicate embryos manifesting poor health (likely due to expression of native genes in the reporter virions). Scale bars represent 50 µm.

**Supplementary Figure S2.**
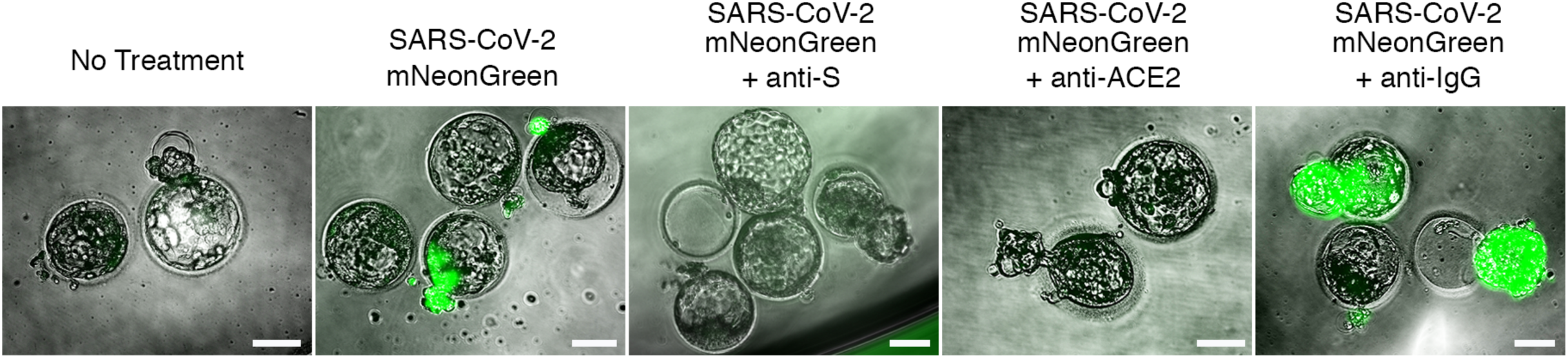
Live SARS-CoV-2 Experiments Indicate Susceptibility to Infection by Cells of the Embryo Through S and ACE2. Sample images from SARS-CoV-2-mNeonGreen experiments, displaying merged brightfield with epifluorescence signal. The sample size of each condition is indicated in Table 2 and Supplementary Table S2. Scale bars represent 100 µm.

